# FateID infers cell fate bias in multipotent progenitors from single-cell RNA-seq data

**DOI:** 10.1101/218115

**Authors:** J.S. Herman, Sagar, D. Grün

## Abstract

Differentiation of multipotent cells is a complex process governed by interactions of thousands of genes subject to substantial expression fluctuations. Resolving cell state heterogeneity arising during this process requires quantification of gene expression within individual cells. However, computational methods linking this heterogeneity to biases towards distinct cell fates are not well established. Here, we perform deep single-cell transcriptome sequencing of ~2,000 bone-marrow derived mouse hematopoietic progenitors enriched for lymphoid lineages. To resolve subtle transcriptome priming indicative of distinct lineage biases, we developed FateID, an iterative supervised learning algorithm for the probabilistic quantification of cell fate bias. FateID delineates domains of fate bias within progenitor populations and permits the derivation of high-resolution differentiation trajectories, revealing a common progenitor population of B cells and plasmacytoid dendritic cells, which we validated by in vitro differentiation assays. We expect that FateID will enhance our understanding of the process of cell fate choice in complex multi-lineage differentiation systems.

## INTRODUCTION

Recent studies utilizing scRNA-seq^1–5^ and single-cell lineage tracing techniques^6–8^, call into question the traditional view of hematopoietic differentiation as a sequence of binary fate choices giving rise to a succession of increasingly fate restricted progenitor types^9^. Evidence from these studies rather suggests early cell fate priming starting at the level of multipotent progenitors (MPP) or even within the HSC pool. Pronounced heterogeneity of common myeloid progenitors (CMP) was elucidated with high resolution^1^, and an early fate bias emerging in human short term HSCs was suggested in a more recent study^2^. However, heterogeneity of lymphoid progenitors has not been well investigated with single-cell resolution. Since lymphoid progenitor heterogeneity was previously found by flow cytometry^10,11^, utilizing distinct sets of cell surface markers in combination with differentiation assays, we here perform an scRNA-seq analysis to comprehensively elucidate heterogeneity across lymphoid progenitors purified from the bone-marrow of adult mice.

Although a number of methods for lineage reconstruction have been developed^12–15^, these algorithms are not specifically designed to uncover subtle transcriptome changes and uniquely assign cells to individual branches without accounting for multi-lineage bias. Weak transcriptome modulations also remain undiscovered by state-of-the-art clustering methods, which partition cells into groups without accounting for the co-existence of fate bias towards multiple lineages within individual cells. To elucidate the process of cell fate emergence, we introduce FateID, a computational method for the quantification of fate bias, manifested by subtle lineage specific transcriptome modulations within a multipotent progenitor population. FateID utilizes prior knowledge to identify committed stages of all lineages and tracks differentiation trajectories backward in time by iteratively applying a random forests-based classification, in order to predict the likelihood of multipotent progenitors to give rise to each lineage. We demonstrate here that FateID reliably detects fate bias in various multi-lineage differentiation systems and, in particular, elucidates lineage choice of lymphoid-biased hematopoietic progenitors.

## RESULT

### Heterogeneity of hematopoietic progenitors in the adult bone marrow

The detection of subtle transcriptome changes reflecting fate biases requires an scRNA-seq method that maximizes both sensitivity and accuracy. Recent benchmarking has demonstrated that CEL-Seq2^16^ optimized both of these factors, but comes with high costs per cell^17,18^. To enable sensitive transcriptome profiling of thousands of cells at low cost, we established an automatized and miniaturized version of the CEL-Seq2 protocol on a mosquito® (HTS version, TTP Labtech) nanoliter-scale liquid handling robot (**Fig. 1a** and Online methods; also see **Supplementary Text 1** and **Supplementary Fig. 1**). Our Mosquito®-CEL-Seq2 (mCEL-Seq2) protocol is compatible with single-cell sorting into 384-well plates and thus allows simultaneous recording of cell surface marker expression by flow cytometry and measurement of single cell transcriptomes by scRNA-seq. To investigate the emergence of cell fate bias towards distinct lymphoid lineage we sequenced altogether 2,880 hematopoietic progenitors sorted from overlapping progenitor populations based on surface expression of Kit and Sca-1 (encoded by *Ly6a*). See **Fig. 1b,c** and **Supplementary Fig. 2**.

**Figure 1.**
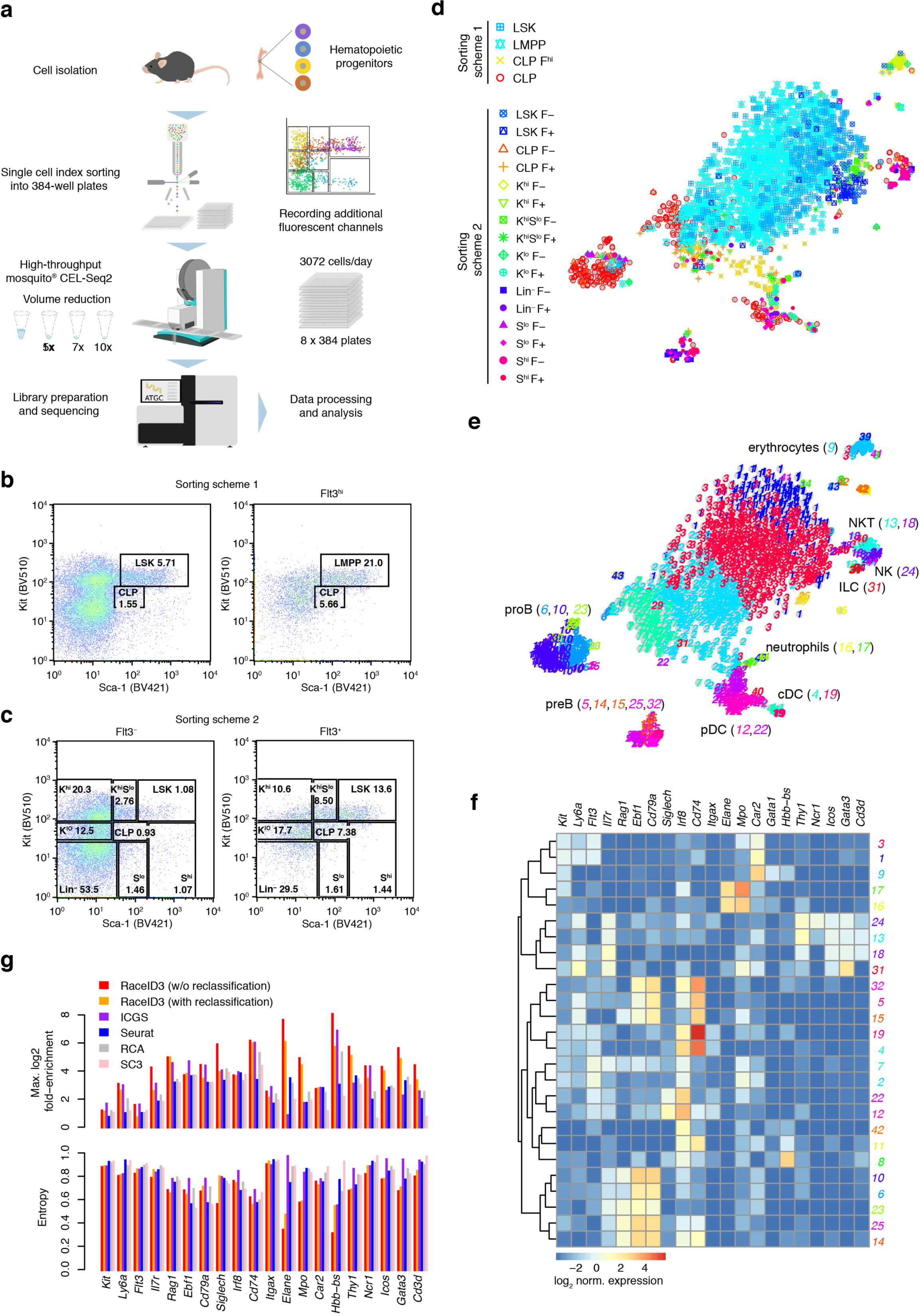
Elucidating transcriptome heterogeneity of multipotent hematopoietic progenitors by single cell RNA-seq. (a) Workflow of robotic CEL-Seq2 implemented on a mosquito® (HTS version, TTP Labtech) nano-liter pipetting robot. The approach enables index-sorting to simultaneously measure single-cell transcriptomes and cell surface marker expression. The robotic approach allows up to 10-fold volume reduction; sensitivity and accuracy were compared to the original CEL-Seq2 protocol for 5-, 7-, and 10-fold reduction (**Supplementary Fig. 1a-e**) (b) The left panel shows the sorting scheme of multipotent Lineage^−^ (Lin^−^) Sca-1^hi^Kit^hi^ (LSK) progenitor cells and Kit^lo^Sca-1^lo^ common lymphoid pogenitors (CLP). The right panel shows the same sorting scheme after gating on Flt3^hi^ cells to enrich for lymphoid-primed multipotent progenitors (LMPP) and Flt3^hi^ CLPs. (c) Sorting scheme for enrichment of Lin^−^ progenitors of all lineages using a tiling window approach on Kit and Sca-1 expression. All populations were divided into Flt3^+^ and Flt3^−^ cells. The percentage of the entire cell population is indicated for each window in (b) and (c). (K: Kit, S: Sca-1) Cells in (b) and (c) were sorted from n=2 mice each. (d) t-SNE map derived by RaceID3 based on transcriptome correlation showing the gate of origin for each cell. Compare legend to (b,c). (e) t-SNE map showing clusters of cells with similar transcriptomes derived by RaceID3. (f) Heatmap of log_2_-transformed averaged normalized expression of known marker genes across clusters. The cluster number and color are indicated on the right. Only clusters with >3 cells were included. A hierarchical clustering dendogram is shown on the right margin. (g) Benchmarking of RaceID3 by comparing to the published Seurat^21^, SC3^23^, RCA^24^, and ICGS^3^ methods. Top: Barplot showing the maximum log_2_-transformed fold-enrichment of a given lineage marker gene across all clusters. Bottom: Barplot showing the entropy of the distribution of average mean expression across clusters for lineage marker genes. To compute the probability used for the entropy calculation, the sum of mean expressions across clusters was normalized to one. Results for the high-resolution settings are shown. See **Supplementary Text 2** and **Supplementary Fig. 5f** for low-resolution settings.

With our sorting strategy we expect to purify cells of all major lineages of the hematopoietic tree (**Fig. 1b,c**). We observed a wide distribution of transcript numbers per cell ranging from a median of 3,034 in common lymphoid progenitors (CLP) to a median of 12,822 in multipotent progenitors (MPP) (**Supplementary Fig. 3a**). A saturation analysis indicated that sequencing was not fully saturated although it started to plateau (**Supplementary Fig. 3b,c**). However, our transcriptome data exhibit high sensitivity compared to several recently published datasets on similar cell populations ^1,2,19^ (**Supplementary Fig. 3d,e**). For further analysis we discarded all cells with <2,000 transcripts, leaving us with 1,949 cells. We analyzed the transcriptome data with our novel RaceID3 algorithm, a substantially improved version of RaceID2^20^ (Online methods).

A t-SNE map of single-cell transcriptome reveals that Lin-Sca^−^1^hi^Kit^hi^ (LSK) cells and lymphoid-primed multipotent progenitors (LMPPs) form a big cloud with CLPs and other small clusters in the periphery (**Fig. 1d**). Cell surface marker expression, quantified from intensities measured by index-sorting for a number of discriminative markers (Kit, Sca-1, Flt3, Il7r, Ly6d) delineates cells from the distinct sorting gates and exhibits moderate correlation with the mRNA level of the respective marker (**Supplementary Fig. 3f-k**).

To resolve heterogeneity within these populations, including rare cell types, we performed clustering analysis with outlier detection using RaceID3. The identified clusters discriminate sub-populations expressing marker genes of distinct lymphoid and myeloid lineages (**Fig. 1e,f** and **Supplementary Fig. 4**). Cluster 6 and 10 comprise *Ebf1*-expressing cells of the B lineage, while cells in cluster 12 express *Siglech*, a marker of plasmacytoid dendritic cells (pDCs). The innate lymphoid branches were resolved into natural killer (NK) cell (cluster 24), NKT cell (clusters 13 and 18) and innate helper lymphoid cell (ILC) type 2 progenitors (cluster 31) based on expression of *Ncr1*, *Cd3d*, and *Gata3*, respectively. We also identified cells of myeloid lineages, in particular, with our alternative sorting strategy (**Fig. 1c**). Clusters 8, 9 and 11 express *Car2*, *Gata1*, and *Hbb-bs* at variable levels and thus represent different stages of erythroblast maturation. Clusters 16 and 17 comprise neutrophil precursors with elevated levels of *Mpo* and *Elane*, while clusters 4 and 19 express *Irf8* and MHCII components and therefore represent conventional dendritic cells (cDC). The central clusters 1, 2, and 3 comprise mainly LSK cells and LMPPs, but already exhibit variable levels of distinct lineage markers. For instance, cluster 1 exhibits up-regulation of *Car2* whereas cluster 3 shows increased expression of *Mpo*. While the former is discriminating the erythroid lineage from neutrophils^1^, the latter has been shown to be expressed in neutrophil and monocyte progenitors^1^. Cluster 7 contains predominantly CLPs and, consistently, exhibits up-regulation of *Il7r* and *Rag1* compared to the other early progenitor clusters 1, 2 and 3.

The performance of RaceID3 on this dataset was benchmarked against a number of alternative methods, i.e. Seurat^21,22^, SC3^23^, RCA^24^, ICGS^3^, based on the expression distribution of known lineage markers across clusters. An ideal clustering method is expected to maximize the fold enrichment of a marker gene in a particular cluster and minimize the spread of the expression domain across clusters and RaceID3 optimizes both metrics (see **Fig. 1g**, **Supplementary Fig. 5** and **Supplementary Text 2**).

In conclusion, we recovered progenitors of the main hematopoietic lineages and found that the pool of early hematopoietic progenitors segregates into sub-populations with transcriptome changes indicative of distinct lineages.

### FateID quantifies fate bias towards distinct lineages in multipotent hematopoietic progenitors

Our clustering analysis suggested population heterogeneity within the multipotent progenitor compartment, correlated with the expression of lineage-specific markers (**Fig. 1f** and **Supplementary Fig. 4**). However, clustering analysis is insensitive to subtle, gradual gene expression changes, potentially indicating the onset of transcriptional priming towards distinct lineages within homogenous multipotent progenitor populations. Moreover, clustering does not estimate the likelihood of a given cell to differentiate towards each lineage. To quantify fate bias of individual progenitor cells from single-cell RNA-seq data, we developed FateID, a supervised algorithm, which starts from committed cell populations of all lineages emerging from a common progenitor, obtained by prior knowledge. FateID iteratively moves backward along the differentiation trajectory to infer transcriptome priming towards each lineage at earlier differentiation stages. More precisely, FateID utilizes random forests^25^, a supervised learning method known for its robustness to over-fitting, to learn the identity of cells in a test set given the training data. FateID starts with target clusters of committed cell populations, identified by clustering or marker gene analysis. Cells in the local neighborhood of each of these target clusters are selected to build a test set. Next, a defined number of cells from the target clusters, most similar to the test set, is used as training set to derive the probability of each cell in the test set to belong to any of the target clusters based on random forests votes. Cells with a significant bias towards a particular lineage become part of the respective target cluster and are utilized for classification in the next iteration. FateID proceeds until the fate biases of all cells have been inferred (**Fig. 2a** and Online methods).

**Figure 2.**
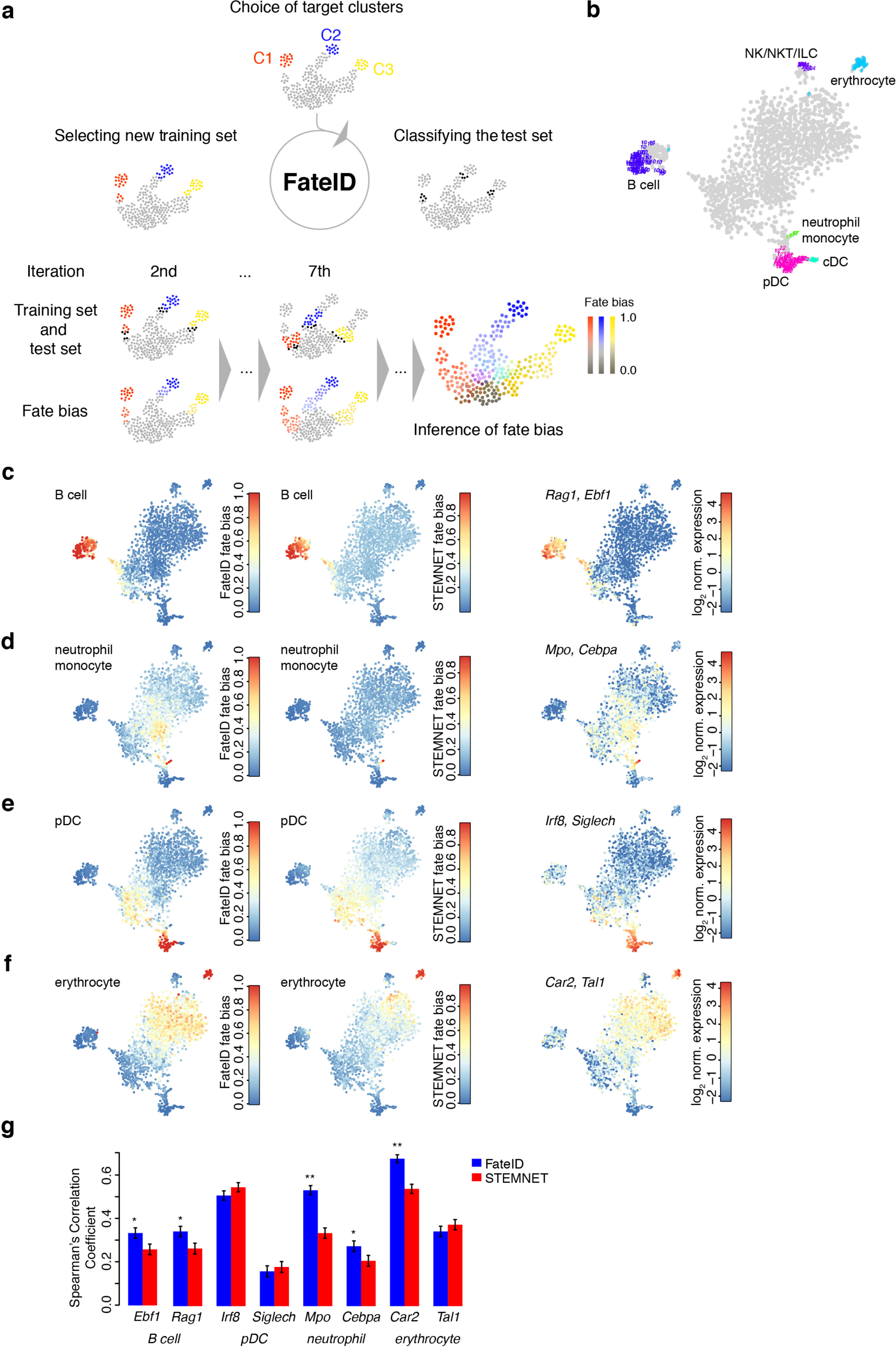
FateID quantifies lineage bias within multipotent progenitor popuations. (a) Pictorial representation of the algorithm. FateID starts with a training set of target clusters of all lineages, comprising more mature stages of differentiation. The algorithm iteratively classifies cells in proximity of the training set (test set) by random forests and thereby expands the training set. To avoid classification of more naïve multipotent cells by genes expressed at more mature stages, the training set in each iteration only comprises cells in the neighborhood of the cells to be classified. See Online methods for details. (b) t-SNE map based on transcriptome correlation after the removal of small outlier clusters and populations of mature and potentially recirculating B cells. The target cluster corresponding to each lineage is highlighted. (c-f) The fate bias, corresponding to the probability of a cell to be assigned to a given lineage, is color-coded in the t-SNE map. The fate bias predicted by FateID (left) and STEMNET (middle) is shown along with log_2_-transformed aggregated normalized expression of two lineage markers. Fate bias and marker gene expression is shown for the B cell (c), the myeloid (d), the pDC (e), and the erythrocyte (f) lineage. (g) Barplot comparing Spearman’s correlation coefficient between the expression levels of lineage markers and fate bias computed by FateID and STEMNET. Error bars and p-values were derived by William’s test statistic (\**P* < 0.05, \*\**P* < 0.001).

We applied FateID to the hematopoietic progenitors with target clusters given by committed states of all lineages as identified by marker genes (**Fig. 1f** and **2b** and Online methods) after removing small outlier clusters with less than five cells. We compared FateID to the recently published STEMNET algorithm^2^. Within the early progenitor clusters comprising LSK cells, LMPPs, and CLPs (clusters 1,2,3,7), FateID could discriminate sub-populations with pronounced fate bias towards B cells, pDCs, neutrophils, and erythrocytes (**Fig. 2c-f**). Neither FateID nor STEMNET could identify substantial fate bias for cDCs and NK/NKT/ILCs (**Supplementary Fig. 6**), potentially due to the lack of intermediate progenitor stages of these lineages in our datasets. This is expected for cDCs, since they terminally differentiate in peripheral organs^26^ and our analysis might have picked up recirculating cells. The fate bias, as quantified by the fraction of random forests votes for the respective target cluster, correlated, within the early progenitor clusters, with the expression of early lineage markers (**Fig. 2c-f** and **Supplementary Fig. 7**), e. g. *Car2* for erythroblasts, *Mpo* for neutrophils and monocytes, *Rag1* for B cells, *Irf8* for pDCs. The correlation derived with FateID was significantly higher than with STEMNET for the majority of genes and comparable for the remaining ones (**Fig. 2g**). Moreover, FateID had in general a larger dynamic range than STEMNET (**Supplementary Fig. 8**). More precisely, while STEMNET divided cells bi-modally into cells with high bias and a finite baseline-level of >10%, the fate bias derived by FateID covers a continuum ranging from zero to one. While STEMNET and FateID could classify more mature stages equally well, STEMNET did not resolve the fate bias in early progenitors as observed by FateID, consistent with expression of early markers. The likely reason for this is that STEMNET classifies progenitors solely based on gene expression in mature cells, while the training set of FateID moves “backward” along the differentiation trajectory with the cells to be classified and hence avoids classifying more naïve cells solely based on markers expressed at mature stages.

Finally, FateID permits the extraction of genes with high importance for classification in any of the iterations (Online methods). Following the relative importance at subsequent iterations, starting from the target clusters, it turns out that markers of more mature stages are initially important, but early stages are classified by different sets of genes (**Fig. 3a** and **Supplementary Fig. 9a-f**), highlighting the relevance of a dynamic training set. For example, classification of the B cell lineage (**Fig. 3a**) starts with cells in cluster 10, highly expressing B cell receptor and surrogate light chain components (*Vpreb1*, *Vpreb3*, *Igll1*). During initial iterations, these genes are most important for classification, while at earlier differentiation stages (later iterations), corresponding to proB cells, *Pax5*, and, even earlier, *Ebf1* become important. Finally, within common lymphoid progenitors, recombination associated genes, such as *Rag1* and *Dntt* become more important for classification. Hence, the feature importance as a function of the differentiation progress reveals stage specific markers and uncovers the complexity of gene regulatory changes during differentiation.

**Figure 3.**
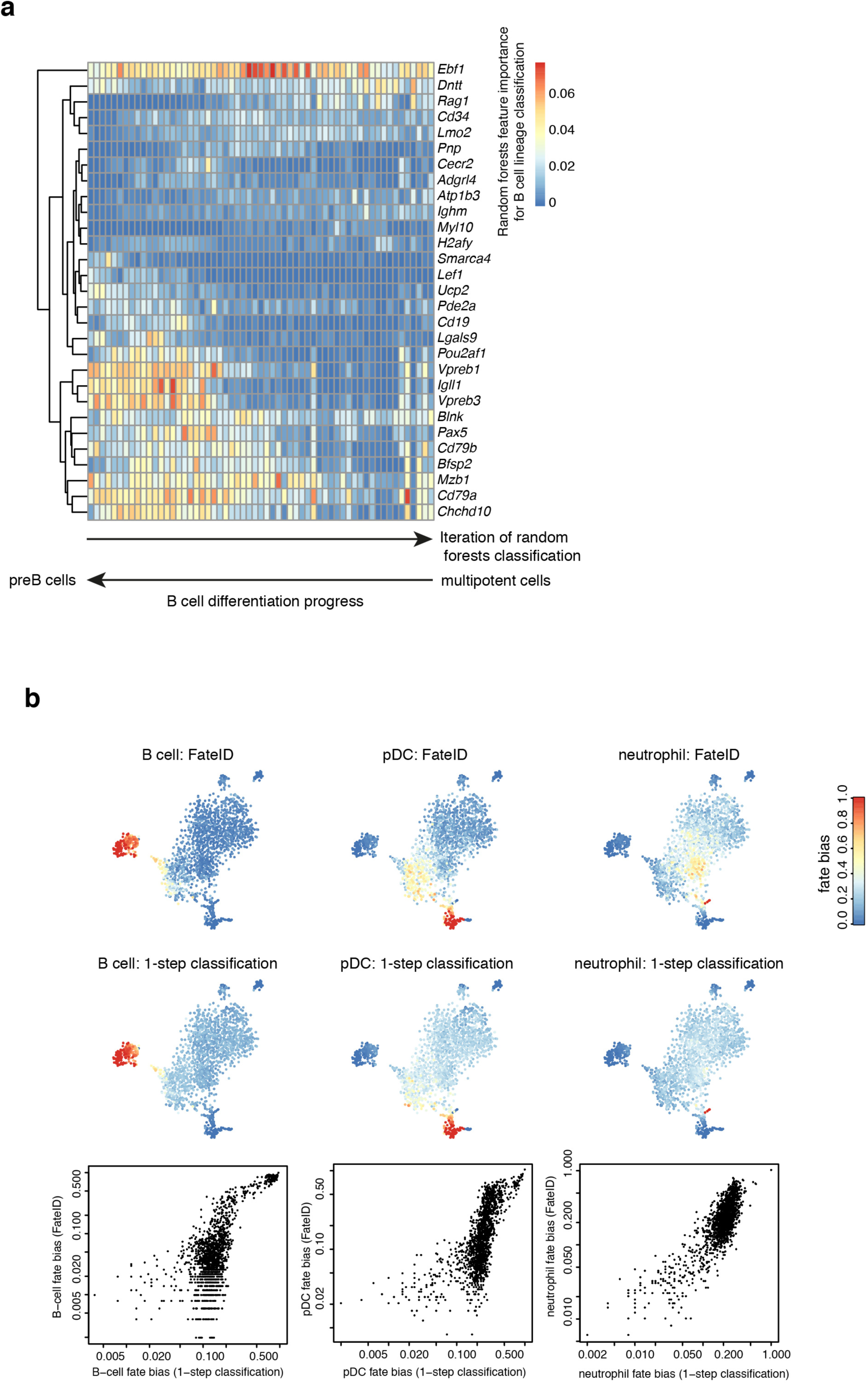
Importance of genes for lineage classification depends on the differentiation stage. (a) Importance of genes during each iteration of the random forests classification of the B cell lineage. The heatmap shows genes with a random forests importance measure >0.02 and a ratio between the absolute importance and its standard deviation >2 for at least a single iteration. Iterations are depicted on the x-axis with the first iteration to the left and the final iteration to the right. Early iterations correspond to more mature stages while late iterations correspond to more naïve stages. A hierarchical clustering dendogram is indicated on the left margin. (b) Comparison of the FateID approach to classification by a single random forests classification of all remaining cells using the target clusters as training set. Examples are shown for the B cell (left), the pDC (center), and the neutrophil (right) lineage. A direct comparison of fate bias estimate for the two approaches reveals saturation at a non-zeo probability in the 1-step classification.

The importance of the iterative classification strategy of FateID is also evident from a comparison to a classification in a single step, where the fate bias distribution across lineages within the multipotent compartment becomes largely uniform (**Fig. 3b**).

In order to enable purification of lineage-biased progenitors we correlated the FateID bias predicted for each lineage to cell surface marker expression measured by index sorting (**Supplementary Fig. 9g**). This analysis confirmed that erythrocyte-biased progenitors localize to a naïve Kit^hi^Sca-1^hi^Flt3^−^Il7r^−^ compartment, while neutrophil bias increases for Kit^hi^Sca-1^lo^Flt3^int^Il7r^−^ cells. Expectedly, B cell lineage bias is enhanced for Il7r^hi^Ly6d^hi^ cells while pDC bias correlates with Flt3 surface expression. Although these observations are consistent with known sorting schemes, additional markers are needed to resolve populations with weak lineage bias.

### FateID reveals a common progenitor of B cells and pDCs

The origin of pDCs and cDCs in the bone marrow is still under debate. While in vitro differentiation assays and genetic lineage tracing suggest a myeloid origin of cDCs^27^, pDCs have mixed origin with a substantial fraction (~26%) exhibiting a history of *Rag1* expression suggesting partial lymphoid origin^28,29^. A comparison of the fate bias for B cells (**Fig. 2c**) and pDCs (**Fig. 2e**) reveals that B cells arise from the *Rag1*^hi^ compartment of the CLP while pDCs are predicted to arise from a progenitor pool encompassing *Il7r*^+^*Rag1*^lo^ CLPs and *Il7r*^−^ LMPPs (**Supplementary Fig. 3**). This observation suggests that the pDC lineage serves as a default pathway of *Il7r*^+^*Flt3*^+^ CLP differentiation unless cells are pushed towards the B cell fate by a secondary signal, e.g. by sufficiently strong IL7 signalling. To further elucidate branching we visualized the cell population in low dimensions using different algorithms provided by FateID (**Fig. 4a** and Online method). A pseudo-temporal order of cells on the B cell, the myeloid, and the pDC differentiation trajectories was inferred by fitting a principal curve to all cells with a fate bias >15% for the respective lineage in a 2-dimensional t-SNE map (**Fig. 4a**). Topological ordering of z-score-transformed pseudo-temporal expression profiles identified stage-specific co-expression modules (**Fig. 4b** and **Supplementary Fig. 10a**). Known regulators of B cell differentiation, e.g., the recombination-activating gene *Rag1*, the preB cell receptor component *Igll1* and proliferation markers such as *Pcna* show the expected pattern reflecting heavy and light chain B cell receptor recombination with intermittent proliferation (**Fig. 4b**). Pseudo-temporal ordering of dendritic cell progenitors recovers a gradual increase of *Irf8*, already during early progenitor stages, and up-regulation of known lineage determining transcription factors, such as *Bst2* and *Tcf4* at later stages (**Supplementary Fig. 10a**). However, early progenitors up-regulate *Il7r* to similar levels like B cell progenitors, but fail to up-regulate *Rag1* to the same extent (**Fig. 4c**). On the other hand, B cell progenitors do not express *Csf1r*, while this receptor is expressed in myeloid progenitors, and, more stochastically, in progenitors with predicted pDC bias (**Supplementary Fig. 10b** and **Fig. 4c**).

**Figure 4.**
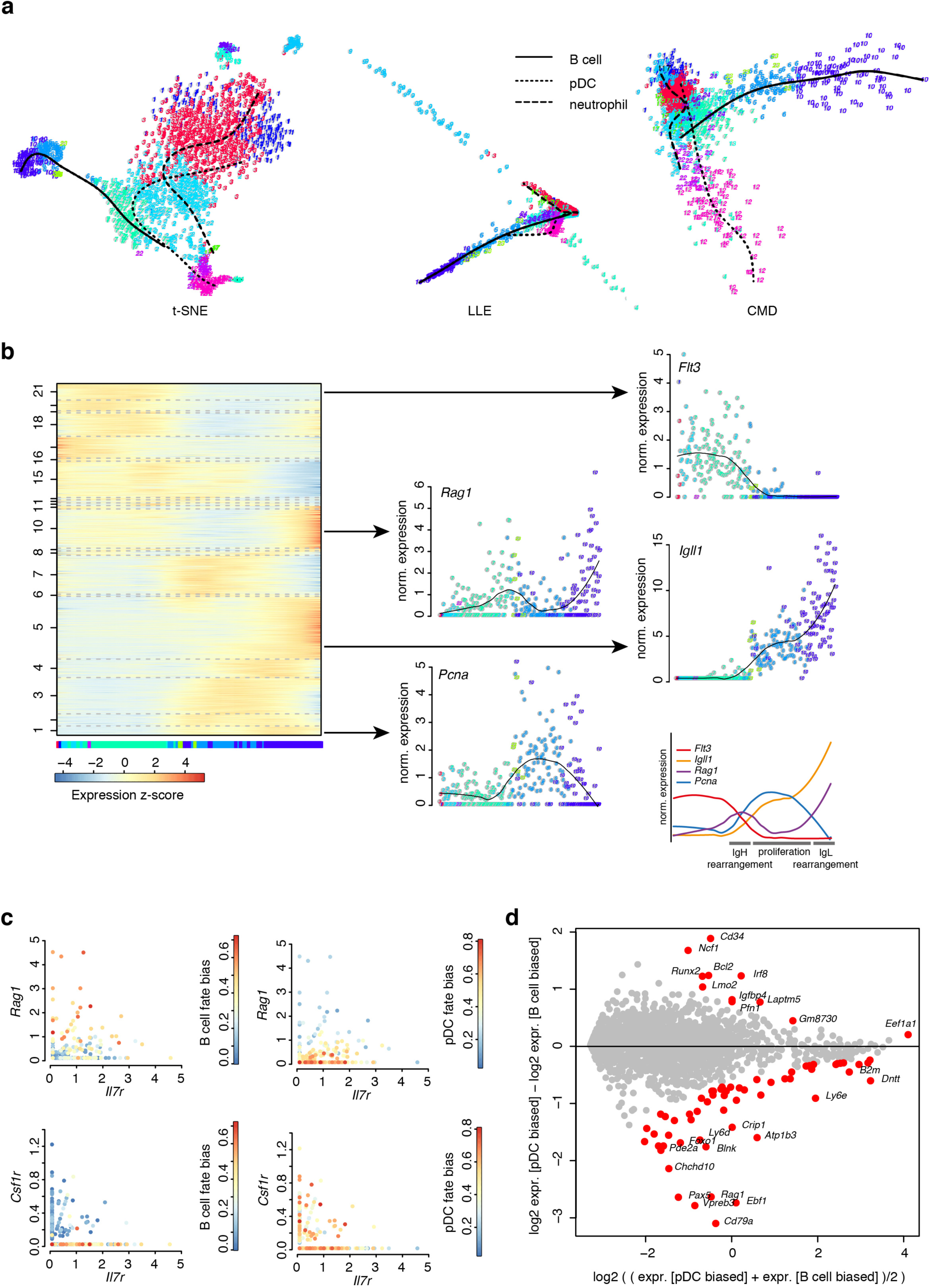
FateID identifies a common progenitor population of B cells and pDCs. (a) Different dimensional reduction representations. Shown are a two-dimensional t-SNE map representation (t-SNE, left), locally linear embedding (LLE, middle), and classical multidimensional scaling representation (CMD, right). Principal curves fitted to cells with a fate bias >0.15 are shown for the B cell, pDC, and myeloid lineages. (b) Self-organizing map of z-score-transformed pseudo-temporal expression profiles along the B-cell developmental trajectory derived from the t-SNE map in (a). Example profiles are shown for four genes dynamically expressed during B cell differentiation. The black line indicates a local regression. The bottom panel shows an overlay of the profiles with known stages of B cell differentiation. (c) Gene-to-gene scatter plots of lymphoid (*Rag1*, *Il7r*) versus myeloid (*Csf1r*) lineage markers, highlighting B cell (left) or pDC (right) lineage bias. (d) MA plot showing differentially expressed genes between cells with larger bias towards the B cell or the pDC lineage. Only cells of cluster 2 and 7 with >0.33 fate bias towards the B cell or pDC lineage were included. Differentially expressed genes with a Benjamini-Hochberg corrected false-discovery rate (FDR) < 0.1 are highlighted in red.

To systematically identify markers of sub-populations with dominating pDC or B cell bias, respectively, within the lymphoid progenitor pool (clusters 2 and 7), we performed a differential gene expression analysis comparing groups of cells with >33% bias towards pDCs and B cells and higher bias towards B cells than pDCs or vice versa (**Fig. 4d**). We found that B cell-biased progenitors already up-regulate known B lineage genes (*Ebf1*, *Pax5*, *Vpreb3*, *Rag1*, *Cd79a*), while higher levels of *Runx2*, *Irf8*, *Ncf1*, *Bcl2*, and *Cd34* were observed in pDC-biased progenitors. The increased importance of these genes during random forests-based classification of the earlier stages is consistent with this observation (**Supplementary Fig. 9b,c**).

Our analysis identifies *Cd34* as a cell surface marker that discriminates between B cell and pDC bias within Il7r^+^Flt3^+^ CLPs (**Supplementary Fig. 10b**). Reassuringly, the previously described marker *Ly6d* for B cell biased CLPs (BLP)^10^ did also come up in our analysis. However, it is also expressed in more mature pDCs (**Supplementary Fig. 10b**) and therefore not ideal to separate B cell and pDC lineage. In conclusion, FateID predicts the existence of a common progenitor of B cells and pDCs and suggests the differentiation towards pDCs as a default pathway in the lymphoid progenitor compartment.

To validate FateID on additional datasets we applied the algorithm to published single-cell transcriptome data for common myeloid progenitors^1^ (see **Supplementary Text 3** and **Supplementary Fig. 11,12**) and intestinal epithelial cells^20^ (see **Supplementary Text 4** and **Supplementary Fig. 13-15**). In contrast to available methods for the inference of lineage trees such as Monocle 2^30^ (**Supplementary Fig. 16**), which perform a fixed assignment of cells to unique branches, FateID uncovers multi-lineage bias within early progenitors that have likely not yet committed to a terminal fate.

### In vitro differentiation assay confirms common progenitor of B cells and pDCs

In order to validate the emergence of pDCs and B cells from a common murine lymphoid progenitor population, we sorted Flt3+ Lin^−^Kit^lo^Sca-1^lo^ cells and sub-gated on Il7r+Cd34-, Il7r+Cd34+, and Il7r-Cd34+ populations (**Fig. 5a** and **Supplementary Fig. 17**). In addition, we sorted LMPPs as in the original experiment. After 7 days of culturing ~400 cells from each of these four populations on OP9 feeder cells in B cell and pDC medium, respectively, we analyzed the relative B cell and pDC lineage output by measuring surface expression of Cd19 and Siglech (see Online methods). In agreement with the FateID predictions (**Fig. 2c,e**), Il7r-Cd34+ positive cells preferentially differentiated into Siglech+ pDCs and Il7r+Cd34+ as well as Il7r+Cd34-cells gave rise to Siglech+ pDC and Cd19+ B cells. We did not observe cells positive for both Cd19 and Siglech. In congruence with the FateID prediction, the fraction of pDCs was markedly reduced by 66% (37%) for Il7r+Cd34-versus Il7r+Cd34+ cells after culturing in B cell (pDC) medium (**Fig. 5b,c**). We confirmed purity of our input populations by single-cell RNA-seq and analyzed these data together with single-cell transcriptome data of the cultured cells (**Fig. 5d**). As expected, the sorted Il7r-Cd34+ and Il7r+Cd34-input populations co-clustered with mixed Flt3+ Lin^−^Kit^lo^Sca-1^lo^ but segregated into sub-populations positive for *Il7r* and *Cd34* mRNA, respectively (**Fig. 5d,e**). In contrast, cultured cells were separated into clusters of B cells, pDCs, cDCs, and lysozyme expressing monocytes (**Fig. 5d,e**). The ratio of the numbers of B cells and pDCs after culturing as determined from single-cell RNA-seq data was in agreement with the flow cytometry data and the FateID predictions, i.e. Il7r-Cd34+ only differentiated into pDCs, while Il7r+Cd34+ and Il7r+Cd34-cells gave rise to both lineages in culture with twice as many B cells versus pDCs for the Il7r+Cd34-cell culture (**Fig. 5f)**. We point out that the sensitivity of the anti-Cd34 antibody is limited and thus discrimination of Cd34+ and Cd34- is not strictly quantitative. Moreover, we observed that B cell clones grow much more rapidly in culture. Therefore, although a strict quantitative comparison is not possible, the assay confirms the FateID prediction of a common lymphoid progenitor population of B cells and pDCs.

**Figure 5.**
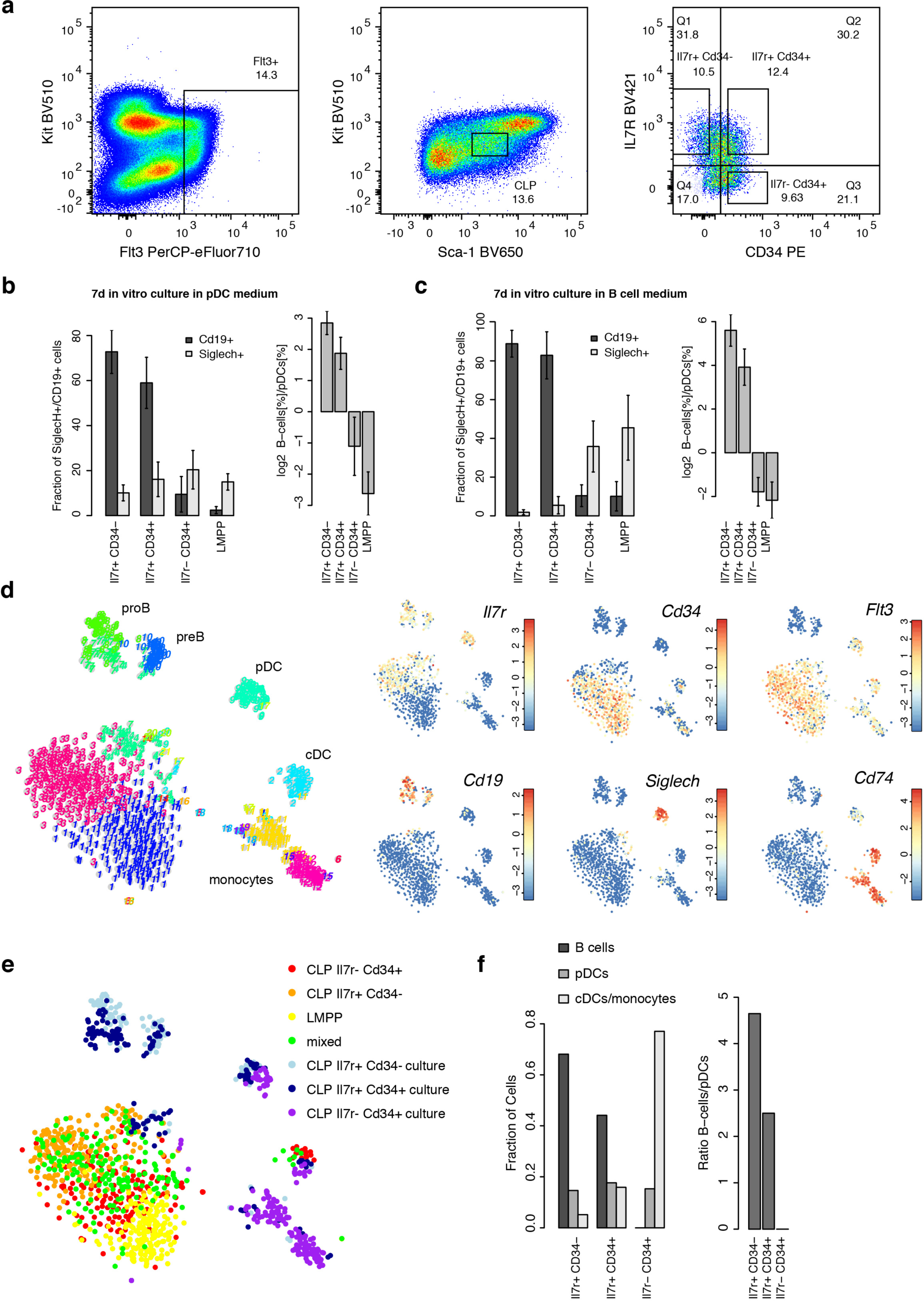
In vitro differentiation assay confirms predicted lymphoid progenitor of pDCs and B cells. (a) Sorting scheme for the purification of Il7r+Cd34-, Il7r+Cd34+, and Il7r-Cd34+ populations from Flt3+ Lin^−^Kit^lo^Sca-1^lo^ cells. The experiment was performed in five biological replicates. Error bars indicate the standard deviation. (b,c) Lineage output of the B cell and the pDC lineage measured by surface expression of Cd19 and Siglech, respectively, after 7 days of culture in pDC (b) and B cell (c) medium for the three sub-populations and LMPPs. The log2-ratio of the fraction of B cells and pDCs is shown to the right. (d) t-SNE map of a RaceID3 analysis of single-cell transcriptomes of cells sequenced for the sub-populations in (a) plus unselected Flt3+ Lin^−^Kit^lo^Sca-1^lo^ cells (mixed) and for cells after culture in pDC medium. Expression of lineage-specific marker genes is shown in the t-SNE maps to the right. (e) t-SNE map highlighting the population of origin. (f) Left: Bar graph showing the relative fraction of cultured cell corresponding to B cells, pDCs, or cDCs/monocytes for the three sub-population sorted from Flt3+ Lin^−^Kit^lo^Sca-1^lo^ shown in (a). Right: Ratio of B-cell and pDC lineage output for same three sub-populations.

### FateID suggests a common progenitor of B cells and pDCs in human

In order to compare the mouse hematopoietic progenitor populations to their counterpart in human, we re-analyzed data from a recent study^2^ (**Supplementary Fig. 18a,b**). As one of the core findings, which was extensively validated by *in vivo* and *in vitro* differentiation experiments, surface expression of CD135 (encoded by *FLT3*) and CD45RA discriminated lymphoid/myeloid- from megakaryocyte/erythrocyte-primed progenitors. FateID analysis confirmed this segregation, but showed a more pronounced separation of the two groups (**Supplementary Fig. 18c)**. Moreover, the correlation of the predicted FateID bias to surface marker expression leads to co-clustering of eosinophil/basophil/mast cell progenitors with megakaryocyte/erythrocyte progenitors. This finding, which is not apparent from the STEMNET predictions, is supported by co-expression of common transcription factors (*GATA2* and *TAL1*) and consistent with a recently described early bifurcation into *Gata1*-positive erythrocyte/megakaryocyte/eosinophil/mast cell progenitors and *Gata1*-negative monocytes/neutrophils/lymphocytes in the murine system^31^.

Moreover, RaceID3 could discriminate two transcriptionally similar lymphoid progenitor clusters, i.e. cluster 3 and 8, which specifically up-regulated genes of the B cell and pDC lineages, respectively, such as *EBF1* and *IRF8*. Both clusters co-express lymphoid genes, such as *DNTT* and *IL7R,* akin to the common progenitor cluster observed in the mouse data. Within this population, FateID discriminated cells with more pronounced bias towards either B cells or pDCs, depending on expression of *IRF8* (**Supplementary Fig. 18d-f**). Finally, FateID predicted increased B cell bias within a more naïve population, coinciding with low expression of the preB cell receptor component *VPREB1*. This remained undetected by STEMNET, which showed a more uniform fate bias in early progenitors (**Supplementary Fig. 18g,h**).

In summary, FateID analysis supports the existence of a common differentiation pathway of B cells and pDCs also in human.

## DISCUSSION

Here, we utilized a down-scaled plate-based implementation of the CEL-Seq2 method, to perform sensitive transcriptome profiling of thousands of lymphoid biased murine bone marrow cells at low cost and developed the FateID algorithm for the estimation of multi-lineage fate biases. We make all our data publicly available on a website (http://hematopoietic-progenitors.ie-freiburg.mpg.de) to facilitate visualization of gene expression domains and fate bias together with differential gene expression analysis.

Our study reveals that the multipotent hematopoietic progenitor population segregates into sub-populations with dominant fate bias towards one of the major lineages, consistent with previous studies suggesting an early fate priming^1,2,6^. A principal curve analysis of the multipotent clusters 1,2,3, and 7 (**Fig. 6a**) suggests successive overlapping expression domains of *Kit*, *Ifitm1*, *Cd34*, *Cd48* and *Flt3* along the pseudo-time axis, concordant with the hypothesized order and mutual similarities of previously described MPP1-MPP4 populations^32,33^ (**Fig. 6b**). This order was also confirmed by StemID2 (**Supplementary Fig. 19**). Our data indicate, consistent with prior findings^34–36^, that cells of megakaryocyte/erythrocyte bias are closest to the MPP1/MPP2 compartment, delineated by up-regulation of *Ifitm1* and *Mki67*^33^, followed by cells of neutrophil/monocyte, pDC, and B cell bias (**Fig. 6c**).

**Figure 6.**
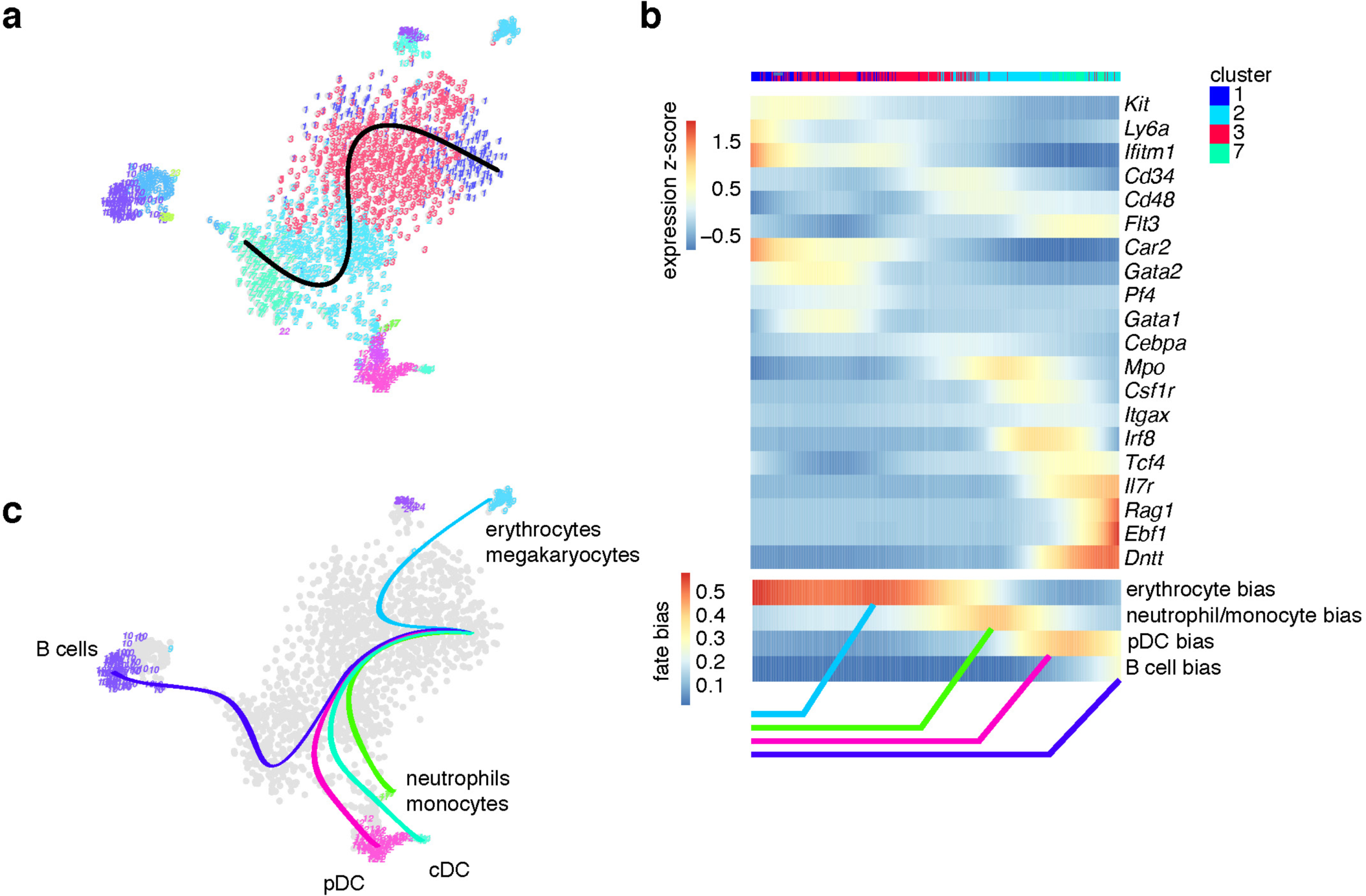
Revised model of hematopoietic differentiation. (a) t-SNE map with a principal curve fitted to all cells within the progenitor clusters (1,2,3,7). (b) Heatmap of z-score transformed pseudo-temporal expression profiles of a number of multipotency and lineage marker genes. Cells were ordered along the principal curve in (a) and profiles were smoothened by a local regression. The bottom panel depicts a local regression of the fate bias using the same temporal ordering. Successive expression of multipotency markers *Kit*, *Ly6a*, *Ifitm1*, *Cd34*, *Cd48*, and *Flt3* is consistent with the ordering of multipotent progenitor (MPP) stages MPP1 to MPP4 as previously inferred by bulk measurements^32,33^. Lineage markers comprise *Gata2*, *Car2*, *Gata1*, *Pf4* for the erythrocyte/megakaryocyte lineage, *Cebpa*, *Csf1r*, *Mpo* for the granulocyte and monocyte lineage, *Itgax* for the conventional dendritic cell lineage, and *Irf8*, *Tcf4* for the plasmacytoid dendritic cell lineage, and *Il7r*, *Rag1*, *Ebf1*, *Dntt* for the B cell lineage. (c) Pictorial representation of the derived lineage tree.

Although our analysis suggests early priming, we could detect domains of overlapping fate bias and validated a common progenitor population for pDCs and B cells: concomitant with up-regulation of lymphoid lineage genes such as *Flt3*, *Dntt* and *Il7r*, cells acquire increased bias towards the pDC lineage. Progenitors with myeloid and pDC biases still express markers of MPPs, such as *Cd34*. Only upon further down-regulation of *Cd34* and concomitant up-regulation of *Il7r* and *Rag1*, cells become more biased towards B cells. A picture emerges where myeloid cells branch off first, followed by pDC progenitors, spanning a continuous spectrum ranging from cells with alternative myeloid potential to cells with alternative lymphoid bias, and an ultimate branching of B cells. This is in line with previous studies suggesting mixed myeloid and lymphoid origin of pDCs^29,37–40^ and could help to resolve this controversy. Hence, our study sheds light on the origin of pDCs in the bone marrow, adding another piece to the puzzle of pDC versus cDC differentiation, which is still subject of intense research^37,38,41–43^.

As another example, we note that FateID recovered a recently characterized bi-potent progenitor of monocytes and neutrophils^3^ (**Supplementary Fig. 20**).

In summary, FateID reveals heterogeneity of multipotent progenitors and links the underlying transcriptome modulations to lineage biases. These predictions shed light on the early regulators of cell fate choice and permit a functional analysis of fate-biased sub-populations if those can be isolated based on *de novo* identified marker genes. We envision that FateID will be a valuable tool for elucidating lineage choice in any multilineage differentiation system.

## Accession Codes

The GEO accession number for the RNA sequencing datasets of murine hematopoietic progenitors reported in this paper is GEO: GSE100037.

## ACKNOWLEDGEMENTS

We would like to thank Andrew Pospisilik for his help in developing mCEL-Seq2. We would like to acknowledge extensive support by Sebastian Hobitz, Konrad Schuldes and Doris Wild for flow cytometry and Ulrike Boenisch for deep sequencing. We would also like to thank Thomas Boehm, Christiane Happe and Rudolf Grosschedl for their valuable input and support. We thank Thomas Boehm, Nina Cabezas-Wallscheid, and Eirini Trompouki for critical reading of the manuscript and valuable feedback.

## AUTHOR CONTRIBUTIONS

J. S. H. performed all experiments, analyzed the data, and created the web-interface. S. established the mCEL-Seq2 protocol with help of J. S. H.. D.G. designed the study, developed the algorithm and wrote the paper. All authors edited the manuscript.

## COMPETING FINANCIAL INTERSTS

The authors declare no competing financial interests.

## ONLINE METHODS

### Mice

5-week old male C57BL/6J wildtype mice were purchased from Charles River or obtained from in-house breedings. Mice were sacrificed in a carbon dioxide chamber and femora and tibiae were collected. All experiments were approved by the Institute’s Review Committee and the local Government.

### Single cell sort of hematopoietic progenitors

Bone marrow of 5-week old male C57BL/6J wildtype mice was isolated from femora and tibiae by flushing using 1x MojoSort™ Buffer (BioLegend®, 480017). Bone marrow cells were filtered (sysmex CellTrics® 30 µm) and cells were counted (Innovatis CASY® Cell Counter). Hematopoietic progenitors were enriched using the MojoSort™ Mouse Hematopoietic Progenitor Cell Isolation Kit (BioLegend®, 480004) according to the manufacturer’s protocol. After the enrichment cells were counted again and 1-5x10^6^ cells per 100 µl 1x MojoSort™ Buffer were stained on ice and in the dark for 20 min using antibodies against c-Kit/CD117 (BV510, BioLegend®, 135119), Sca-1 (BV421, BioLegend®, 108127), Flt3/CD135 (PerCP-eFluor® 710, eBioscience™, 46-1351-80), CD48 (APC-Cy7, BioLegend®, 103432) or Ly-6C (APC/Fire™ 750, BioLegend®, 128045), CD27 (PE, BD, 558754) or SiglecH (PE, BioLegend®, 129605), Ly-6D (Alexa Fluor® 647, BD, 561147), IL-7Rα/CD127 (PE/Dazzle™ 594, BioLegend®, 135031) and the lineage markers CD3e (FITC, BD, 553062), Ter119 (FITC, eBioscience™, 11-5921-82), B220 (FITC, BD, 553088), Gr-1/Ly-6G (FITC, BD, 553127), CD11b/Mac-1 (FITC, eBioscience™, 11-0112-85). The cells were sorted on a BD Influx™ cell sorter using single cell mode and index information was recorded. Cells were sorted into 384-well plates containing 240 nl lysis buffer (see below) and 1.2 µl of hydrophobic encapsulation barrier (Vapor-Lock, Qiagen, 981611). After the single cell sort, the plates were centrifuged at 2200 g at 4˚C for 10 min, snap-frozen thereafter with liquid nitrogen and stored at −80˚C until they were processed.

### Amplified RNA preparation from single cells

The CEL-Seq2 protocol, as previously described^16^, was adapted for the use with a nanoliter pipetting robot (mosquito® HTS, TTP Labtech). The Volumes used in the original protocol were reduced by 5-fold for the single cell sort of hematopoietic progenitors and all the reagent ratios were kept the same. Hematopoietic progenitors were sorted into 384-well plates (Corning, PCR-384-RGD-C). Every well was filled with 240 nl lysis buffer. Lysis buffer was prepared using the following reagents with final ratio in parentheses: 10 mM dNTP (1/12), 1:100000 ERCC mix 1 or 2 (2/12), water with 0.35% Triton® X-100 (7/12) and 25ng/µl of 1 out of 192 uniquely barcoded polydT primers with UMI (2/12). 1.2 µl of hydrophobic encapsulation barrier (Vapor-Lock, Qiagen, 981611) were added onto the lysis buffer in every well. The sorted plates were incubated at 90˚C for 3 min and cooled to 4˚C thereafter. For cDNA first strand synthesis 160 nl of First Strand Reaction Mix containing SuperScript™ II Reverse Transcriptase and First Strand Synthesis Buffer (Invitrogen, 18064014) and RnaseOUT™ (Invitrogen, 10777019) was added and the plates were incubated at 42˚C for 1 hour followed by heat-inactivation at 70˚C for 10 min. Second strand synthesis was performed at 16˚C for 2 h by adding 2196 nl of Second Strand Reaction Mix containing E. coli DNA Polymerase I (Invitrogen, 18010025), E. coli DNA Ligase (Invitrogen, 18052019), RNaseH (Invitrogen, 18021071) and Second Strand Buffer (Invitrogen, 10812014). For cDNA purification, 96 or 192 wells containing uniquely barcoded single cell samples were pooled and 0.8 volumes of Agencourt® AMPure XP beads (Beckman Coulter, A63881) were mixed with the pooled samples and incubated for 15 min at room temperature. The samples were put on a magnetic stand (96 well format) and incubated for 5min, the supernatant was removed and the beads were washed twice by adding 180 µl of 80% ethanol and by incubating for 40 s. The beads were dried for 10 min and resuspended in 7 µl nuclease free water (Invitrogen™, AM9937). For the generation of amplified RNA (aRNA) 1.6 µl from each nucleotide solution (ATP, CTP, GTP, UTP), 1.6 µl of 10x Reaction Buffer and 1.6 µl of T7 Enzyme from the MEGAscript® T7 Transcription Kit (Invitrogen, AM1334) were added to the eluted cDNA and incubated at 37˚C for 13 h. After in vitro transcription 6 µl of Exo-SAP-IT™ Express PCR Product Cleanup Reagent (Applied Biosystems™, 75001.1.ML) was added and the samples were incubated for another 15 min at 37˚C. Thereafter, aRNA was fragmented by adding 2.44 µl 10x RNA Fragmentation Buffer and incubating at 94 ˚C for 3 min and fragmentation was stopped by adding 2.44 µl RNA Fragmentation Stop Solution (NEB, E6150S). For aRNA purification, 0.8 volumes of Agencourt® RNAclean XP beads (Beckman Coulter, A63987) were added to the sample and incubated for 15 min at room temperature. After that, samples were placed on a magnetic stand and incubated for 5 min, the supernatant was removed and the beads were washed three times by adding 180 µl 70% ethanol and incubating for 40 s. Ethanol was removed, beads were air-dried for 10 min and resuspended in 7 µl of nuclease free water.

### Single cell library preparation from aRNA

The aRNA was reversely transcribed in order to generate libraries for sequencing, as previously described^16^. 1 µl of custom random hexamer RT primer (GCCTTGGCACCCGAGAATTCCANNNNNN) and 0.5 µl of 10mM dNTP were added to the aRNA. The samples were incubated at 65˚C for 5 min and cooled to 4˚C afterwards. Thereafter 4 µl of First Strand Reaction Mix containing SuperScript™ II Reverse Transcriptase and Buffer (Invitrogen, 18064014) and RnaseOUT™ (Invitrogen, 10777019) was added and samples were first incubated at 25˚C for 10 min followed by incubation at 42˚C for 1 hour. To every sample 2 µl of one uniquely indexed RPI index primer and 2 µl of RP1 primer (TruSeq Library Prep, sequences available from Illumina), 25 µl of Phusion® High-Fidelity PCR Master Mix with HF Buffer (NEB, M0531) and 11 µl of nuclease-free water were added. PCR was performed using 98˚C for 30 s as initial step, followed by 11 cycles of amplification (98˚C for 10 s, 60˚C for 30 s, 72˚C for 30 s) and a final extension at 72˚C for 10 min. To each sample 1 volume (50 µl) of Agencourt® AMPure XP beads (Beckman Coulter, A63881) was added and the samples were incubated at 25˚C for 15 min. The samples were incubated for 5 min on a magnetic stand (96 well format), the supernatant was removed and the beads were washed twice by adding 180 µl of 80 % ethanol and by incubating for 40 s. Ethanol was removed and beads were dried for 10 min at room temperature. DNA was eluted with 25 µl nuclease-free water and the purification was repeated with an adjusted volume of magnetic beads (25 µl). After the final step the samples were eluted in 10 µl nuclease-free water. DNA concentration and fragment size were determined using the Qubit® dsDNA HS Assay Kit (Invitrogen, Q32854) and the Agilent High Sensitivity DNA Kit for Bioanalyzer (Agilent Technologies, 5067-4627), respectively. Libraries were sequenced on an Illumina HiSeq 2500 System in high output run mode at a depth of ~200,000 reads per cell.

### Mouse embryonic stem cell culture

Mouse embryonic stem cells (mESC) were cultured in KnockOut™ DMEM (Gibco, 10829018) supplemented with 15% KnockOut™ Serum Replacement (Gibco, 10828028), 1x non-essential amino acids (Gibco, 11140050), 1mM sodium pyruvate (Gibco, 11360070), 1x GlutaMAX™ (Gibco, 35050061), 1x Penicillin/Streptomycin (Gibco, 10378016), 5µg/ml Insulin (Sigma, I0516), 100 µM 2-mercaptoethanol (Gibco, 31350010) and 1000 U/ml LIF (Millipore, ESG1107). Medium was changed on daily basis. Confluent cells were de-attached by incubating with Trypsin-EDTA (Gibco, 25200056) at 37˚C for 5 min. mESC were split every 3 - 5 days with 1:30 - 1:50 split ratios.

### Single cell library preparation for different volume reductions

The ratios of reagents from the original CEL-Seq2 protocol were maintained for all the volume reductions. The protocol for different volume reductions was conducted as described above except that the volumes for the lysis buffer plate, first strand synthesis and second strand synthesis were adjusted. For every condition 48 wells of a 384-well plate were prepared with 1.2 µl, 240 nl, 171 nl or 120 nl lysis buffer corresponding to 1-fold (original CEL-Seq2 volume), 5-fold, 7-fold or 10-fold reductions, respectively. Every well contained 20 nl of 1:75000 diluted ERCC mix 1. Lysis buffer was covered with hydrophobic encapsulation barrier (Vapor-Lock, Qiagen, 981611). mESC were harvested using Trypsin-EDTA (see above) and single cells were sorted into each well using the BD Influx™ cell sorter. For first strand synthesis the adjusted volumes 0.8 µl, 160 nl, 114 nl or 80 nl of First Strand Reaction Mix were used for 1-fold, 5-fold, 7-fold or 10-fold volume reductions, respectively. Second strand synthesis was performed using 11 µl, 2196 nl, 1569 nl or 792 nl of Second Strand Reaction Mix for the different volume reductions, respectively. The rest of the protocol was performed as described above.

### Staining and sorting of hematopoietic progenitors for cell culture and single cell sequencing

Bone marrow of 5-week old male C57BL/6J wild type mice was isolated from ilia, femora and tibiae by crushing using 1x MojoSort™ Buffer (BioLegend®, 480017). Bone marrow cells were filtered (sysmex CellTrics® 30 µm) and cells were counted (Innovatis CASY® Cell Counter). Hematopoietic progenitors were enriched using the MojoSort™ Mouse Hematopoietic Progenitor Cell Isolation Kit (BioLegend®, 480004) according to the manufacturer’s protocol. After the enrichment cells were counted again and 5-7x10^6^ cells per 100 µl 1x MojoSort™ Buffer were stained on ice and in the dark for 40 minutes using antibodies against c-Kit/CD117 (BV510, BioLegend®, 135119), Sca-1 (BV650, BioLegend®, 108143), Flt3/CD135 (PerCP-eFluor® 710, eBioscience™, 46-1351-80), CD48 (APC/Fire™ 750, BioLegend®, 103445), CD34 (PE, BD, 551387), Ly-6D (Alexa Fluor® 647, BD, 561147), IL-7Rα/CD127 (BV421, BD, 566377) and the lineage markers CD3e (FITC, BD, 553062), Ter119 (FITC, eBioscience™, 11-5921-82), B220 (FITC, BD, 553088), Gr-1/Ly-6G (FITC, BD, 553127), CD11b/Mac-1 (FITC, eBioscience™, 11-0112-85), CD19 (FITC, BioLegend®, 101505), Siglec-H (FITC, BioLegend®, 129603). The cells were sorted on a BD FACSAria™ Fusion cell sorter. For cell culture, cells from different gates (see sorting strategy) were sorted using 4-way purity mode into 100 µl RPMI 1640 medium (complete formulation, Thermo Fisher Scientific, A1049101) supplemented with 10% Fetal Calf Serum (Corning BV, 35-016-CV), 1x Penicillin/Streptomycin (Gibco, 10378016) and 55µM 2-Mercaptoethanol (Gibco, 21985023). For single cell sequencing, single cells were sorted using single cell mode and index information was recorded. Cells were sorted into 384-well plates containing 240 nl lysis buffer (see below) and 1.2 µl of hydrophobic encapsulation barrier (Mineral Oil, Sigma-Aldrich, M8410). After the single cell sort, the plates were centrifuged at 2200 g at 4˚C for 10 minutes, snap-frozen thereafter with liquid nitrogen and stored at −80˚C until they were processed using the down-scaled version of CEL-Seq2 (mCEL-Seq2).

### Cell culture of sorted hematopoietic progenitors

Sorted hematopoietic progenitors were seeded onto Mitomycin C-treated OP9 feeder cells. Approximately 400 cells were plated in one well of a 96-well plate and grown in 200 µl of medium at 37 ˚C and 5% CO_2_. For the differentiation of the sorted progenitors towards plasmacytoid dendritic cells RPMI 1640 medium (complete formulation, Thermo Fisher Scientific, A1049101) was used and supplemented with 10% Fetal Calf Serum (Corning BV, 35-016-CV), 1x Penicillin/Streptomycin (Gibco, 10378016), 55µM 2-Mercaptoethanol (Gibco, 21985023) and 200 ng/ml FLT3L (BioLegend, 550704).

In order to differentiate the sorted cells towards the B cell lineage Opti-MEM (Thermo Fisher Scientific, 31985062) was used and supplemented with 10% Fetal Calf Serum (Corning BV, 35-016-CV), 1x Penicillin/Streptomycin (Gibco, 10378016), 55 µM 2-Mercaptoethanol (Gibco, 21985023) and 5 ng/ml IL-7 (BioLegend®, 577802), 10 ng/ml SCF (BioLegend®, 579702) as well as 10 ng/ml FLT3L (BioLegend®, 550704). On the following day after sorting 100 µl of medium was removed and replaced with fresh medium. Every two days 100 µl of medium was replaced with fresh medium. The cells were grown for 7 days and thereafter harvested to perform staining on lineage markers and to sort them for single cell sequencing.

### Staining of cultured hematopoietic progenitors for flow cytometry analysis and single cell sequencing

Cultured cells were detached by resuspension and washed once with 1x MojoSort™ Buffer (BioLegend®, 480017). Staining of the cells was performed in 50 µl 1x MojoSort™ Buffer for 40 minutes using antibodies against c-Kit/CD117 (BV510, BioLegend®, 135119), Sca-1 (BV650, BioLegend®, 108143), Flt3/CD135 (PerCP-eFluor® 710, eBioscience™, 46-1351-80), CD34 (PE, BD, 551387), Ly-6D (Alexa Fluor® 647, BD, 561147), IL-7Rα/CD127 (BV421, BD, 566377), Siglec-H (FITC, BioLegend®, 129603) and CD19 (APC/Fire™ 750, BioLegend®, 115557). Cells were washed with 1x MojoSort™ Buffer and resuspended in 1x MojoSort™ Buffer again. Stained cells were analyzed on a BD LSRFortessa cell analyzer or sorted using the BD FACSAria™ Fusion cell sorter into 384-well plates containing 240 nl lysis buffer (see below) and 1.2 µl of hydrophobic encapsulation barrier (Mineral Oil, Sigma-Aldrich, M8410) for single cell library preparation using CEL-Seq2.

### Preparation of OP9 feeder cell layer

OP9 feeder cells were grown in 1x MEM Alpha (Gibco, 12561-056) supplemented with 20% Fetal Calf Serum (Corning BV, 35-016-CV), 1x Penicillin/Streptomycin (Gibco, 10378016) and 55 µM 2-Mercaptoethanol (Gibco, 21985023). OP9 cells were grown until 80 - 90% confluency and detached using Trypsin-EDTA (Gibco, 25200056) at 37˚C for 5 minutes. Cells were resuspended in fresh complete MEM Alpha medium and cell concentration was adjusted to seed approximately 40000 - 50000 cells in 100 µl of medium per well of a 96-well plate.

### Quantification of transcript abundance

Paired end reads were aligned to the transcriptome using bwa (version 0.6.2-r126) with default parameters^44^. The transcriptome contained all gene models based on the mouse ENCODE VM9 release downloaded from the UCSC genome browser comprising 57,207 isoforms derived from 57,207 gene loci with 57,114 isoforms mapping to fully annotated chromosomes (1 to 19, X, Y, M). All isoforms of the same gene were merged to a single gene locus and gene loci were merged to larger gene groups, if loci overlapped by >75%. This procedure resulted in 34,111 gene groups. The right mate of each read pair was mapped to the ensemble of all gene groups and to the set of 92 ERCC spike-ins in sense direction. Reads mapping to multiple loci were discarded. The left mate contains the barcode information: the first six bases corresponded to the cell specific barcode followed by six bases representing the unique molecular identifier (UMI). The remainder of the left read contains a polyT stretch and adjacent gene sequence. The left read was not used for quantification. For each cell barcode and gene locus, the number of UMIs was aggregated and, based on binomial statistics, converted into transcript counts^45^.

### RaceID3

RaceID3 is an improved version of the previously published RaceID2 algorithm^20,46^. In comparison to RaceID2 a number of additional optional steps has been implemented. A major change is the availability of a feature selection step. If feature selection is enabled, genes will be selected with a variability exceeding the baseline variability as inferred by a second order polynomial fit of the expression variance of all genes as a function of the mean after log-transformation. Only these genes will be used for the computation of the distance matrix used in the k-medoids clustering. For the outlier identification step, all genes remaining after expression filtering are still used to rescue the identification of cell population that were potentially missed due to the feature selection step. A second major change is the introduction of a random forest based reclassification: after outlier identification the robustness of the resulting partition is tested by random forest classification^25^ using the entire dataset as test set with out-of-bag sampling. The final cluster membership is subsequently determined as the cluster with the highest number of random forest votes.

RaceID3 furthermore offers a number of normalization methods. The normalization by rescaling to the median transcript count has been replaced by rescaling to the minimum total transcript number of all cells that survived the filtering, since the previous version led to inflated transcript counts in some cells and, subsequently, to false positive outlier cells. This method is now the default mode and preferable over down-sampling, since the latter comes with a loss of information. In addition, RaceID3 offers alternative normalization schemes, which are described in detail in the reference manual.

The improved algorithm further permits the elimination of unwanted variability. As a minimally invasive approach signature genes of unwanted sources of variability, such as cell cycle or experimental batch, can be provided and all genes with significantly correlated expression to one of these signature genes are removed for clustering and outlier identification.

As a second option, a PCA-based approach has been implemented. Expression data are log-transformed prior to PCA and genes in the tail of the loadings distribution for each principal component are screened for an over-representation of defined sets of signature genes (e. g. annotated cell cycle genes). If a significant overrepresentation is detected, the respective principal component is removed prior to back-transformation.

A third method eliminates unwanted sources of variability by a linear regression. This method is akin to the limma method^47^ and particularly useful for the correction of batch effects.

To identify signature genes for each cluster, RaceID3 now infers differentially expressed genes by utilizing an approach similar to DESeq^48^, but with a dispersion parameter of the negative binomials inferred from the internal background model.

Finally, we could substantially improve the run time compared to the previous version.

The RaceID3 and StemID2 algorithms are available at github together with a detailed reference manual: https://github.com/dgrun/RaceID3_StemID2

### StemID2

The logic of StemID2 remains unchanged compared to the original StemID algorithm. However, an alternative method for the inference of significant links of the lineage tree has been implemented to circumvent to time-intense randomizations of the lineage tree and allow for running StemID on datasets with thousands of cells. In this approach a large randomized distance matrix (with a number of cells equal to the size of the entire dataset) is created only once for each cluster *i* and, keeping all cluster medoids the same as in the non-randomized version, randomized cells are projected onto links between the medoid of cluster *i* and all clusters *j ≠ i*. The average population frequency of each link in the randomized state is now inferred from these projections and significant overrepresentation of cells on a link in the original data inferred by a binomial test requiring a minimum p-value. Although this approach is much faster it only provides an approximation of the actual significance, since the correlation of the population between links is neglected.

Functions for follow-up analysis of gene expression dynamics have also been integrated in StemID3. Pseudo-temporally ordered cells on a differentiation trajectory defined as a succession of significant links can be extracted to group the pseudo-temporal gene expression profiles into co-expression modules by using self-organizing maps. A similar approach is implemented in the FateID algorithm and described in the method section of this algorithm.

The RaceID3 and StemID2 algorithms are available at github together with a detailed reference manual: https://github.com/dgrun/RaceID3_StemID2

### RaceID3 analysis of lymphoid cells, myeloid cells, and intestinal cells

Prior to processing of mouse lymphoid pogenitors by RaceID3, cells with high expression (>2% of all transcripts) of *Kcnq1ot1*, a previously identified marker of low quality cells^20^, were removed. Moreover, transcript correlating to *Kcnq1ot1* or any of the genes *Gm10715*, *Gm42418*, *Gm10800* with a Pearson’s correlation coefficient >0.65 were also removed. Finally, all reads mapping to ERCC spike-ins were discarded. For the analysis of lymphoid progenitors, RaceID3 was run with mintotal=2000, minexpr=3, outminc=3, FSelect=TRUE, probthr=10^−4^ and random forests-based reclassification. For the benchmarking, RaceID3 was also run without random forests-based reclassification to produce a high resolution clustering with a larger number of outliers. In both cases, we initialized CGenes with the following set of genes in order to remove cell cycle and batch associated variability: *Pcna*, *Mki67*, *Ptma*, *Hsp90ab1*, *Actb*, *Jun*, *Fos*, *Gnas*, *Hspa8*. Batch signature genes were identified by a differential gene expression analysis between cells of different batches. FGenes was initialized with *Malat1* and *Igkc*, since these genes were highly expressed across a number of unrelated cell types. For the myeloid progenitor data, we used the published quantification^1^ and RaceID3 was run after downsampling with mintotal=3000, CGenes=*Hsp90ab1*, minexpr=5, outminc=5, FSelect=TRUE, probthr=10^−4^ and random forests-based reclassification. For the intestinal data, we used the published quantification^20^ and RaceID3 was run with mintotal=2000, minexpr=3, outminc=3, FSelect=TRUE, probthr=10^−3^ and random forests-based reclassification. For human progenitors we applied RaceID3 to the published quantification^2^ with mintotal=10000, minexpr=167, outminc=167, FSelect=TRUE, probthr=10^−3^. However, distances were computed with metric=”spearman”, i.e. Spearman’s correlation was used instead of Pearson’s correlation. This alleviated the effect of enhanced variability of the expression values, which correspond to normalized read counts instead of UMI counts for this dataset. Based on the saturation behavior of the within-cluster dispersion a cluster number of 14 was chosen. CGenes was initialized with the following set of genes in order to remove cell cycle associated variability: *PCNA, MKI67, MCM5, MCM7*. On mouse progenitor data from Olsson et al.^3^, RaceID3 was run with mintotal=3000, minexpr=5, outminc=5, FSelect=TRUE, probthr=10^−4^ and random forests-based reclassification.

### Clustering methods for benchmarking of RaceID3

The Seurat algorithm^22^ was run with the same parameters as in the online tutorial. Based on the PC elbow plot the first 13 principal components were used for clustering. A high-resolution clustering with resolution=2 and a low-resolution clustering with resolution=1 was performed. The RCA^24^ method was run in the SelfProjection mode after changing the upper_thresh parameter in the featureConstruct function to upper_thresh = max(log10(5), median(x[x > min_val])), since the original value was too large for UMI-based data. Clustering was executed in a high-resolution mode (deepSplit_wgcna=2, min_group_Size_wgcna=3) and a low-resolution mode (deepSplit_wgcna=1, min_group_Size_wgcna=10). SC3^23^ was run with 10 or 20 clusters, to obtain low-resolution and high-resolution clustering. ICGS^49^ was run on non-normalized transcript counts with quantile normalization and default parameters otherwise.

### FateID

The FateID algorithm performs an iterative calculation in order to infer fate bias in multipotent progenitors, starting from cells within committed states of the distinct lineages arising from a common progenitor. These states are addressed as target states and it is expected that cells within each of these states do not exhibit mixed lineage signatures anymore. To run FateID, an expression matrix of all genes in all cells and a partitioning of cells into cell states or types are required. These cell states are expected to comprise all target states and at least one common state for all the remaining cells. However, finer partitions of the entire cell pool are also permitted, e. g. as obtained from a clustering algorithm. In this case, the set of target states has to be defined as a subset of the partition. FateID also offers the possibility to infer target states based on the expression of a list of marker genes, with an arbitrary number of markers for each lineage. In this mode, FateID will compute the aggregated mean expression of the markers in the top *n* cells, and select all cells with an aggregated marker gene expression greater than this threshold. This selection step is optionally followed by an initial random forests reclassification to test if any of the remaining cells can be classified with high confidence as one of the target states. This step can also be used to perform feature selection by importance sampling (see explanation below).

Starting from the target states, FateID will proceed backward in differentiation time to iteratively learn progenitors of the target states by using random forests classification^25^. Briefly, random forests is an ensemble learning method constructing a multitude of decision trees during learning and returning the mode of the classes for each instance across all trees. Due to the inherent bootstrapping strategy (bagging) combined with random selection of features, random forests are robust against overfitting. The only adjustable parameter to which the results of random forests are somewhat sensitive is the number of features *m* sampled at each node. This number is kept constant across all decision trees and was also kept constant across FateID iterations. In this implementation it was set to the default value √*M* where *M* is the number of features (i.e. genes).

The FateID algorithm utilizes random forests in the following way: in a given iteration, the training variables are the cells within the target states with transcript counts for all expressed genes as features. A prior feature selection for the entire data set, e. g., by extracting strongly variable genes, might enhance the performance of the algorithm. The response variable is the partition of the target states. Now, a fixed number of *m* cells in the neighborhood of each of the *k* target states are selected. For target state *i*, these cells are the *m* cells with the smallest median distance to the cells in state *i*, as measured by 1-Pearson’s correlation of the transcript count vectors of two cells. Alternatively, arbitrary distance matrices can be provided for this selection step. The set of *k* x *m* cells represents the test set for the random forest iteration. For the random forest classification, the training set can be reduced to the number of *h* cells for each target state with the shortest distance to any of the cells in the test set selected for this target state. After classification, cells from the test set with a significant fate bias towards a given target state *i*, i. e. significantly more votes for state *i* than for any other state (with *P<0.05* based on random counting statistics) are assigned to this target state for the next iteration. The algorithm proceeds this way until all cells have been tested.

By this strategy, cells populating early stages of differentiation are not classified solely based on markers expressed in mature stages, but rather based on markers of immediate subsequent stages of differentiation.

The number *m* of cells to be selected for testing in each round and the number *h* of cells from each target state are the two main parameters of the algorithms and should be chosen dependent on the coverage of the dataset. Larger values of *h* provide higher confidence of the classification but also diminish the influence of the local neighborhood. Large datasets with sufficient coverage of all differentiation stages permit larger values of *h*.

The test set size *m* should be selected based on similar consideration: large values decrease the running time, but could lead to a decrease in specificity when early progenitors are tested against distant mature stages. In practice, with datasets of hundreds to few thousands of cells presented in this manuscript we use *m=5* and *h=20*, but FateID was found to be very robust to changes in these parameters.

*Extracting feature importance from FateID.* The features that are relevant for classification within each random forest iteration are informative and comprise marker genes either enriched or depleted during progenitor stages of a given target state. The random forest classification is performed with importance sampling and thus returns for each gene the mean decrease in classification accuracy upon permutation of this gene within each out-of-bag sample, and its respective standard deviation. FateID extracts all genes with importance greater than a defined threshold and an importance z-score (mean importance/standard deviation of importance) exceeding another defined threshold. These genes are grouped by hierarchical clustering and displayed as a function of the iteration number, starting at the initial target state and ending at the most naïve state. This strategy of interrogating the data reveals the role of different gene sets during subsequent stages of differentiation.

The fate bias for each cell is quantified by the fraction of votes for each of the target states and can thus be interpreted as a probability. We note that this probability is likely to be highly correlated to the actual probability of differentiating into the corresponding lineage, but will not be identical to this value, since external signals might modify a pre-existing bias.

The FateID algorithm is available as an R package from github with an extensive vignette: https://github.com/dgrun/FateID.

### Running FateID and STEMNET on hematopoietic progenitors, myeloid progenitors, and intestinal cells

For the mouse hematopoietic progenitors and the myeloid progenitors, FateID was run with minnr=5 and minnrh=20. For the intestinal data and the human hematopoietic progenitors, we chose minnr=10, since the size of the dataset was smaller, comprising only few cells for some of the lineages. Intestinal target clusters were inferred by FateID, using the following list of marker genes: *Alpi* (enterocytes), *Clca3* (goblet cells), *Lyz1* (Paneth cells), *Dclk1* (tuft cells), *Chgb* (enteroendocrine cells). Target clusters for human hematopoietic cells were inferred by FateID, using the following list of marker genes: *EBF1* (B cells), *IRF8* (pDCs), *LYZ* (monocytes), *ELANE* (neutrophils), *LMO4* (eosinophils/basophils/mast cells), *HBB* (erythrocytes), *ITGA2B* (megakaryocytes). For the mouse hematopoietic progenitors and the myeloid progenitors we selected the respective RaceID3 clusters as target clusters. STEMNET^2^ was initialized with the same target clusters as FateID and run with default parameters. For data from Olsson et al.^3^ target clusters were inferred by FateID, using the following list of markers: *Pf4* (megakaryocytes), *Epor* (erythrocytes), *Cebpe* (neutrophils), *Ly86* (monocytes); FateID was run with minnr=5 and minnrh=20.

### Visualization of fate bias

FateID allows visualization of the data structure by a number of dimensional reduction representations, i. e. t-SNE^50^, classical multidimensional scaling, locally linear embedding, and diffusion maps^12^. These representations can be computed for dimensional reduction to an arbitrary number of dimensions *k*. For *k=2* or *k=3*, these representations can be inspected directly, while for higher dimensions FateID permits visualization after projecting on a subset of dimensions. Within these representations FateID can highlight the partition, the expression of genes, or the fate bias to facilitate inspection of the results.

### Inference of pseudo-temporal order and co-expression modules

To inspect gene expression dynamics during differentiation, FateID performs a topological ordering of pseudo-temporal expression profiles by self-organizing maps (SOM) as utilized previously^20^. To obtain a pseudo-temporal order, all cells with a significant fate bias towards a given unique target state, or, alternatively, all cells with a fate bias towards this state larger than a defined threshold, are selected. These cells can be pseudo-temporally ordered by FateID using principal curve analysis within a selected dimensional reduction representation, which best reflects the differentiation dynamics indicated by a gradient in the fate bias. The weight of the cells in the initial target cluster is increased tenfold to enforce that the principal curve traverses this set of cells, and an optional starting cluster can be provided to initialize the inference with a principal curve connecting this cluster with the initial target state. Alternatively, other methods such as diffusion pseudotime^12^ can be used. FateID has implemented a direct interface to diffusion pseudotime ordering. To identify modules of co-expressed genes along a differentiation trajectory to a defined target states the expression levels in the pseudo-temporally ordered cells are smoothened by local regression after z-score transformation. These pseudo-temporal gene expression profiles are topologically ordered by computing a one-dimensional self-organizing map (SOM) with 1,000 nodes. Due to the large number of nodes relative to the number of clustered profiles, similar profiles are assigned to the same node. Only nodes with more than 5 assigned profiles are retained for visualization of co-expressed gene modules. Neighboring nodes with average profiles exhibiting a Pearson’s correlation coefficient >0.9 are merged to common gene expression modules. These modules are depicted in a final map.

### Differential gene expression analysis

Differentially expressed genes between two subgroups of cells were identified similar to a previously published method^48^. First, we infer a negative binomial distribution reflecting the gene expression variability within each subgroup based on a background model for the expected transcript count variability computed by the same strategy as in RaceID2^20^. Using these distributions a p-value for the observed difference in transcript counts between the two subgroups is computed as described DESeq^48^. These p-values were corrected for multiple testing by the Benjamini-Hochberg method. We implemented a function to perform this calculation, with the alternative option to output the results of a DESeq2^51^ analysis.

### Running Monocle2 on lymphoid progenitors

Monocle 2^30^ was run on non-normalized transcript counts with min_expr=3 and num_cells_expressed >= 5. Cell types of the B cell, pDC, erythrocyte, neutrophil, cDC, and innate lymphoid lineage were assigned based on expression of Ebf1, Siglech, Gata1, Elane, Cd74, and Thy1 require a minimum expression of 5 for all cell types but cDC. For cDC, at least 50 transcripts of Cd74 were required. For subsequent clustering, differentially expressed genes (qval < 0.01) were identified for each cell types and the top 100 genes for each cell type were selected. Clustering was performed with 8 clusters. Dimensions were reduced using the same set of genes as used for clustering with max_components=2, norm_method = 'log'.

### Data availability and software

Primary read data and processed count files for the single-cell RNA-seq datasets of murine hematopoietic reported in this paper can be downloaded from GEO (accession number GSE100037). We make all our processed expression data publicly available on a website (http://hematopoietic-progenitors.ie-freiburg.mpg.de) to facilitate visualization of gene expression domains and fate bias together with differential gene expression analysis.

Accession codes for the myeloid progenitor^1^ and the intestinal epithelial^20^ single-cell transcriptome data can be found in the referenced studies. RaceID3, StemID2 and are available on the github repository: https://github.com/dgrun/RaceID3_StemID2.

The FateID algorithm is available as an R package from github with an extensive vignette: https://github.com/dgrun/FateID.

